# Vaccinia virus vaccination is expected to elicit highly cross-reactive immunity to the 2022 monkeypox virus

**DOI:** 10.1101/2022.06.23.497143

**Authors:** Syed Faraz Ahmed, Muhammad Saqib Sohail, Ahmed Abdul Quadeer, Matthew R. McKay

## Abstract

Starting May 2022, a novel cluster of monkeypox virus infections was detected in humans. This has spread rapidly to non-endemic countries and sparked global concern. Vaccinia virus vaccines have demonstrated high efficacy against monkeypox viruses in the past and are considered an important outbreak control measure. Viruses observed in the current outbreak carry distinct genetic variation that have the potential to affect vaccine-induced immune recognition. Here, by investigating genetic variation with respect to orthologous immunogenic vaccinia-virus proteins, we report data that anticipates vaccine-induced immune responses to remain highly cross-reactive against the newly observed monkeypox viruses.

## INTRODUCTION

The monkeypox virus outbreak being observed in 2022 (MPXV-2022) is highly distinctive. Unlike previous outbreaks of monkeypox, which were localized [1–3] and resulted in small numbers of infections due to limited ability for human-human transmission [4–6], the emerging outbreak has already resulted in over 3300 reported cases spanning more than 40 countries [7], in just a few weeks since the first case was reported on May 7, 2022 [8]. This has raised significant public health concerns and has prompted authorities to coordinate intervention and control strategies. The underlying determinants of the MPXV-2022 outbreak remain unclear [9], however phylogenetic analyses [10] of genomic sequences reported on GISAID [11] from at least 15 countries place these in the West African clade of MPXV (MPXV-WA). This is surprising given the historically low outbreak-causing potential observed for this particular clade [12–14].

Vaccination based on replication competent or highly attenuated vaccinia virus (VACV) is seen as a primary strategy for combating monkeypox outbreaks in humans. Studies on human monkeypox disease outbreaks in the 1980s had shown that a replication competent VACV vaccine, used for the eradication of smallpox [15], offered 85% protection against the Congo Basin clade of MPXV (MPXV-CB) [4,16]. However, there is still a lack of scientific data on vaccine effectiveness in humans against viruses belonging to the MPXV-WA clade, which appears most relevant to the sequences observed in the current outbreak [10].

With the aim of investigating expected cross-reactive immunity provided by VACV vaccines against the MPXV-2022 outbreak, we used available genomic sequence and immunological data to quantify genetic similarities and differences between the VACV immunogenic proteins and their orthologs in MPXV-2022 isolates. VACV vaccines are known to elicit both antibody (humoral) and T-cell (cellular) responses in humans [17–21]; therefore, we analyzed VACV proteins and epitopes known to be specific targets of antibodies and T cells.

While identifying a small number of mutations of potential immunological consequence, our data broadly indicates that MPXV-2022 is highly genetically conserved within immunogenic protein regions and epitopes of VACV, and it will likely exhibit similar cross-reactive humoral and cellular immunity profiles upon VACV vaccination as for MPXV-CB. The high reported efficacy of VACV vaccines against MPXV-CB may, therefore, be similarly anticipated for MPXV-2022, though this cannot be established without clinical evaluation data.

## METHODS

### Acquisition and pre-processing of sequence data

A total of 75 complete genome sequences of MPXV-2022 were downloaded on June 7, 2022, from the GISAID database (https://www.gisaid.org/) (Supplementary Table 1). The complete genome reference sequences for the VACV and MPXV-CB were downloaded from NCBI using the GenBank accession IDs NC_006998 and NC_003310 (Zaire-96-I-16), respectively. Similar to our previous works [22,23], an in-house bioinformatics pipeline was developed to align these nucleotide sequences and then translate them into amino acid residues according to the coding sequence positions provided along the reference sequence for VACV and the location of the gene (whether on the forward or reverse strand).

The MAFFT software was used to perform all multiple sequence alignments [24]. A total of 182 genes with coding sequences are described along the VACV reference sequence and the corresponding translated genomic regions of MPXV-2022 sequences represent the 182 MPXV-2022 ortholog proteins. Coding sequence of one VACV protein, C3L, did not align with valid coding regions within the genomic sequences of MPXV-2022, possibly due to partial gene deletion and/or poor quality of sequence alignment within these regions. Therefore, C3L was excluded from the analysis.

### Computing genetic similarity

Genetic similarity between any pair of nucleotide or protein sequences was computed from their pairwise sequence alignments. All positions within the pairwise alignment that had a gap were counted as insertion/deletion (indel). For nucleotide pairwise alignments, all positions where there was a mismatch of nucleotides were counted as single nucleotide polymorphisms (SNPs), while for protein pairwise alignments, all positions where there was a mismatch of residues were counted as substitutions. Genetic similarity was defined as the fraction of positions in the pairwise alignments that had no SNPs/substitutions or indels.

### Acquisition of epitope data

VACV-derived B cell and T cell epitopes were searched on the Immune Epitope Database (IEDB) (https://www.iedb.org/; accessed 10 June 2022) [25] by querying for the virus species name: “Vaccinia virus” (taxonomy ID: 10245) from “human” hosts. The search was limited to include only experimentally determined epitopes [26] that were associated with at least one positive assay: (i) Positive B cell assays for B cell epitopes, and (ii) positive T cell assays for T cell epitopes. For the B cell epitopes, only two epitopes were found (IEDB IDs: 735903 and 735904), both within the D8L protein. A literature search identified one additional VACV antibody epitope within D8L [27] that is reported to be targeted in humans. For the T cell epitopes, the search was restricted to epitopes with lengths between 9 and 21 residues, which covers the typical range of both HLA class I and class II epitopes. This search returned 388 T cell epitopes in total.

### Selection of immunogenic proteins

The eight VACV proteins identified as targets of neutralizing antibodies in humans (A17L, A27L, A28L, A33R, B5R, D8L, L1R, and H3) have been widely reported in the literature [28,29]. The 121 VACV proteins identified as targets of T cells were obtained by mapping the 388 VACV-derived T cell epitope sequences onto the VACV protein sequences.

### Visualization of protein crystal structures

VACV protein crystal structures were obtained from the Protein Databank (www.rcsb.org) and the structural figures were made using the PyMOL software (www.pymol.org).

## RESULTS

The newly identified MPXV-2022 sequences demonstrate a mean genetic similarity of ∼84% with the VACV reference sequence (GenBank: NC_006998.1). There exist ∼3% SNPs and ∼13% indels, which translates to ∼6.5k SNPs and ∼27.5k indels due to the large genome sizes of MPXV-2022 and VACV (∼200kbp). Investigation of the genetic differences within immunogenic proteins that are targets of either B cells/antibodies or T cells can offer insights into the anticipated effects of this genetic variation on immune recognition by VACV vaccine-induced responses.

### VACV proteins targeted by neutralizing antibodies share high sequence similarity with MPXV-2022 orthologs

Eight VACV immunogenic proteins are known to elicit neutralizing antibodies (NAbs) [28,29]. A subset of these have been used as antigens in subunit vaccines against smallpox and monkeypox [30–35]. The similarity of the eight proteins with respect to their orthologs in MPXV-2022 sequences and the MPXV-CB reference sequence was evaluated. The latter serves as a meaningful reference since the efficacy of VACV vaccines against monkeypox outbreaks caused by MPXV-CB has been reported previously [4,16]. For all eight immunogenic proteins, there was a high genetic similarity (range: ∼94% to ∼98%) between VACV and both the MPXV-2022 consensus sequence and the MPXV-CB reference sequence (Table 1). The high genetic similarity observed for the MPXV-2022 consensus sequence was also observed when considering the complete set of available MPXV-2022 sequences (Supplementary Fig. 1).

**Table 1.**
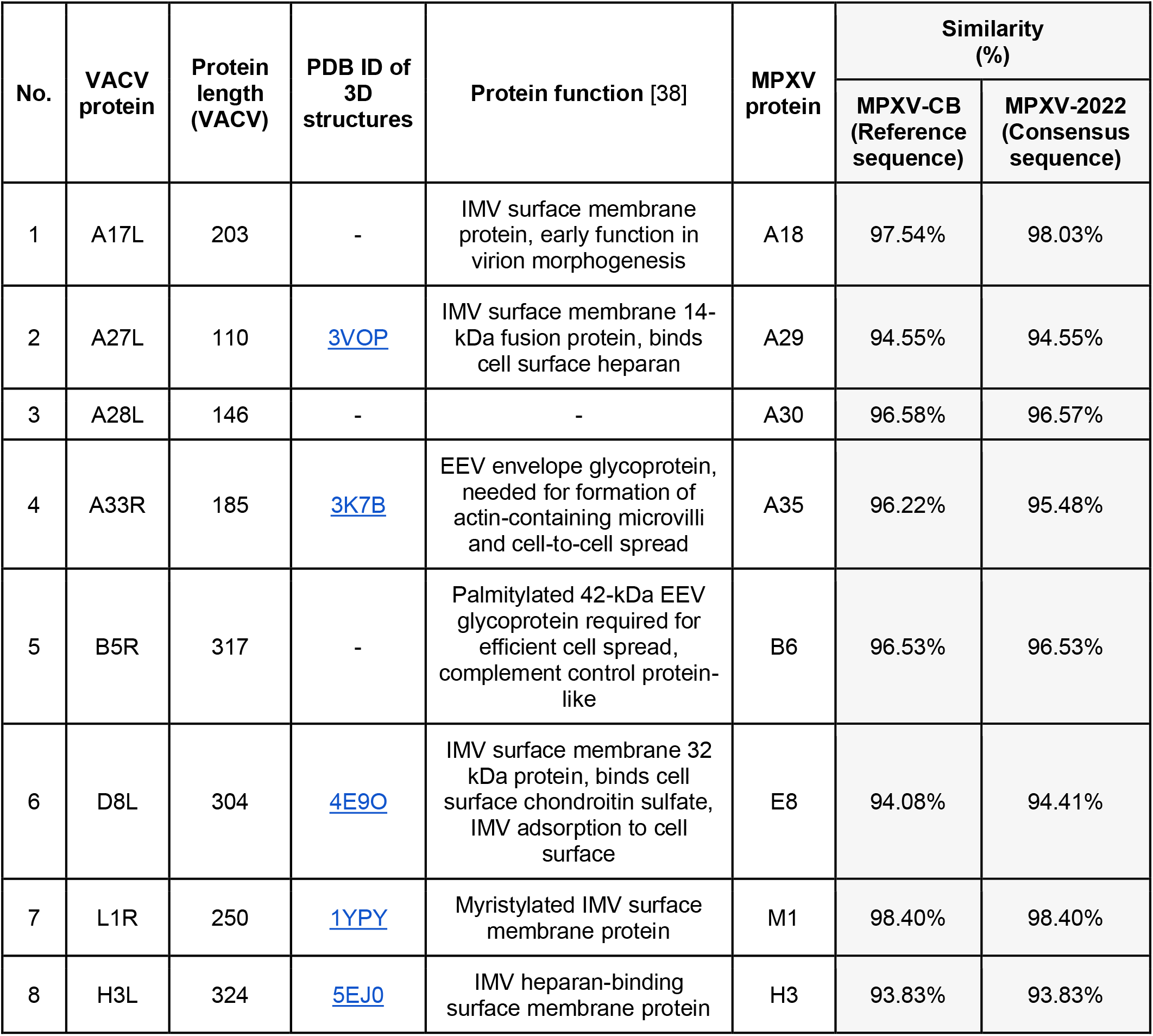
Summary of the VACV proteins known to be targets of NAbs.

Further examination of the specific mutations between the immunogenic proteins of VACV and both the MPXV-2022 consensus sequence and the MPXV-CB reference sequence revealed exactly the same sets of mutations for half (4/8) of the proteins. For the remaining proteins, most mutations were still in common. In these cases, the MPXV-2022 consensus or the MPXV-CB reference sequence carried one or two additional mutations (Table 2). The highest number of common mutations relative to VACV were observed for the H3L and D8L proteins (19 and 17 mutations respectively). Mapping these mutations onto the reported crystal structures (H3L: [PDB ID 5EJ0] and D8L: [PDB ID 4E9O]) revealed that, for H3L, 7/19 common mutations appear to be exposed (Fig. 1A). The unique mutations in MPXV-2022 (A4V) and MPXV-CB (T111I) are also seemingly exposed (Fig. 1A); hence, they may be accessible to targeting antibodies and have the potential to affect binding or neutralization. For D8L, 7/17 mutations common in MPXV-2022 and MPXV-CB, as well as the unique mutation in MPXV-CB (A19T), are all exposed (Fig. 1B). A subset (4/7) of the common mutations and the unique mutation in MPXV-CB overlap with the known binding footprints [27] of three D8L-specific VACV NAbs (Fig. 2). The potential impact of these mutations on antibody binding and neutralizing capacity remains to be determined. Importantly, for all three antibodies, MPXV-2022 had no additional mutations in the binding footprints relative to MPXV-CB. Overall, while some genetic differences were observed between the immunogenic proteins of VACV targeted by NAbs and their orthologs in MPXV-2022, most of these differences were common to MPXV-2022 and MPXV-CB. VACV-vaccine-induced humoral immunity against MPXV2002 is therefore anticipated to be similar to that against MPXV-CB.

**Table 2.** Genetic comparison of known VACV NAb target proteins with corresponding proteins in MPXV-CB and MPXV-2022. The substitutions are computed with reference to the VACV proteins and using the consensus sequence of the ortholog protein for MPXV-2022.

| VACV Protein | Length | MPXV | Substitutions |  |
| --- | --- | --- | --- | --- |
| A17L | 203 | MPXV-CB | T183I | Y155F, R165K, T171P, V188I |
|  |  | MPXV-2022 |  |  |
| A27L | 110 | MPXV-CB | K27N, A30T, D39Y, E40G, V61I, R74H |  |
|  |  | MPXV-2022 |  |  |
| A28L | 146 | MPXV-CB | G23S, Q110R, V130I, V131A, A137T |  |
|  |  | MPXV-2022 |  |  |
| A33R | 177 | MPXV-CB | L59Q | A73S, Q117K, L118S, S120E, T127A, I141T |
|  |  | MPXV-2022 | E67K, A88V |  |
| B5R | 317 | MPXV-CB | Q50S, S55L, I82V, N87D, A166V, M188I, V233I, I236T, T240S, V283M, V296I |  |
|  |  | MPXV-2022 |  |  |
| D8L | 304 | MPXV-CB | A19T | N18D, P23T, S65T, L66I, L114I, S118A, L124S, T146M, K163T, V181A, D209E, A246V, R253K, T261A, E272G, F293L, R296Q |
|  |  | MPXV-2022 |  |  |
| L1R | 250 | MPXV-CB | L51I, K177R, V242I, M248I |  |
|  |  | MPXV-2022 |  |  |
| H3L | 324 | MPXV-CB | T111I | L17P, P44Q, N48D, V51I, K53N, A63V, N74D, N108K, K117N, V124I, M143I, M168I, N171D, L230M, A233S, N251T, A263V, T265A, A274T |
|  |  | MPXV-2022 | A4V |  |

**Figure 1.**
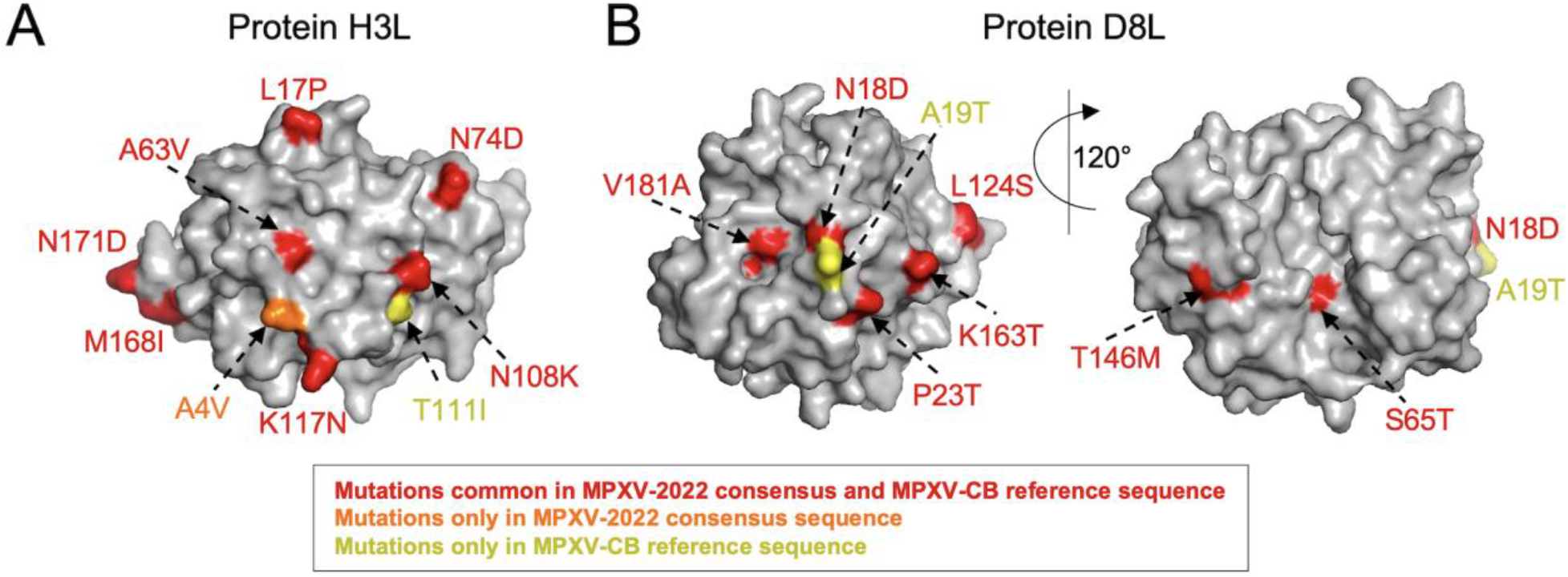
Mapping mutations observed in MPXV-2022 and MPXV-CB on the structure available for VACV (**A**) H3L [PDB ID: 5EJ0] and (**B**) D8L [PDB ID: 4E9O] surface proteins. The core structure of each protein is shown in gray, while mutations and their labels are colored according to the scheme in the legend.

**Figure 2.**
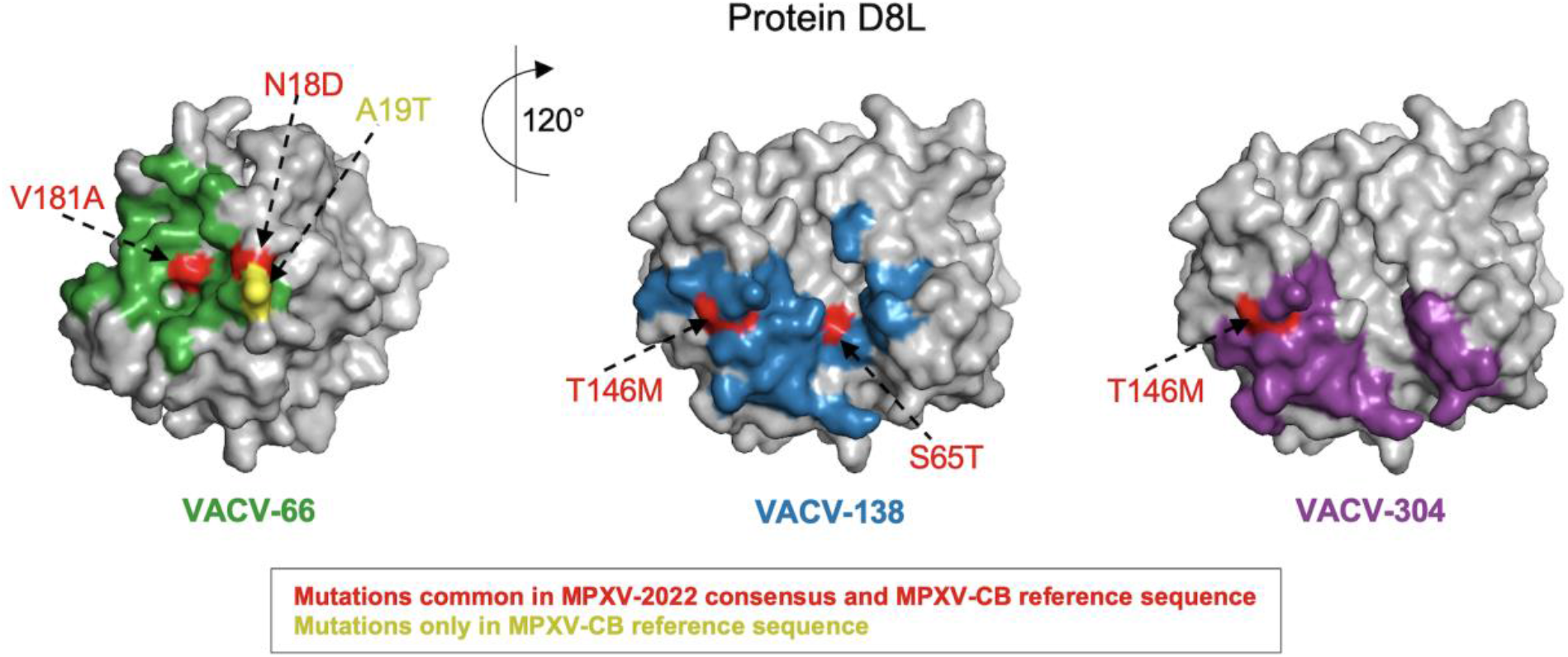
MPXV-2022 does not comprise any new mutation relative to the MPXV-CB reference sequence in the epitopes of the known D8L-specific antibodies. Mutations observed in MPXV-2022 and MPXV-CB are mapped on the epitope of the three known neutralizing antibodies (VACV-66, VACV-138, and VACV-304) in the D8L protein [PDB ID: 4E9O]. The structure of D8L is shown in gray, while mutations and their labels are colored according to the scheme in the legend.

### VACV proteins and epitopes targeted by T cells are largely conserved in MPXV-2022

Experimental studies have investigated T cell responses induced by VACV vaccines [17,20]. We collated a set of all VACV proteins associated with at least one reported T cell epitope (Methods), revealing a total of 121 proteins. A high degree of genetic similarity was observed between these VACV proteins and their orthologs in MPXV-2022 and MPXV-CB (Supplementary Fig. 2). This would anticipate significant cross-reactivity of VACV-induced T cell responses against MPXV-2022 (and MPXV-CB).

Across these 121 proteins, 388 VACV-derived T cell epitopes (197 CD8^+^ and 191 CD4^+^) were identified. Comparing the epitope sequences with the MPXV-2022 consensus and the MPXV-CB reference sequence revealed that 71.6% (278/388) of the epitopes had an exact match in both MPXV-2022 and MPXV-CB. Of the remaining, 1.55% (6/388) differed only in MPXV-2022, 1.8% (7/388) differed only in MPXV-CB, while 25% (97/388) differed in both MPXV-2022 and MPXV-CB (Fig. 3 and Supplementary Table 2). That is, despite high genetic similarity between VACV proteins and the MPXV-2022 and MPXV-CB orthologs (Supplementary Fig. 2), genetic variation was observed in over one quarter of the T cell epitopes. The potential impact of this variation on T cell recognition remains to be determined. Importantly, further examination of the 25% of VACV T cell epitopes that differed in both MPXV-2022 and MPXV-CB showed that a large fraction (70%) associated with epitope mutations were identical in both MPXV-2022 and MPXV-CB (Fig. 3). Taken together, sequences of 89.2% of the T cell epitopes are identical in both MPXV-2022 and MPXV-CB. Interestingly, for the eight immunogenic proteins targeted by NAbs, a high percentage (93%) of the known VACV T cell epitopes had an exact match in MPXV-2022. Our analysis, overall, shows that most known VACV T cell epitopes are fully conserved in MPXV-2022, and that while notable variation within VACV T cell epitopes still exists in MPXV-2022, most of this variation is identical to that observed for MPXV-CB. Hence, VACV-vaccine-induced cellular immunity against MPXV-2022 is anticipated to be similar to that observed for MPXV-CB.

**Figure 3.**
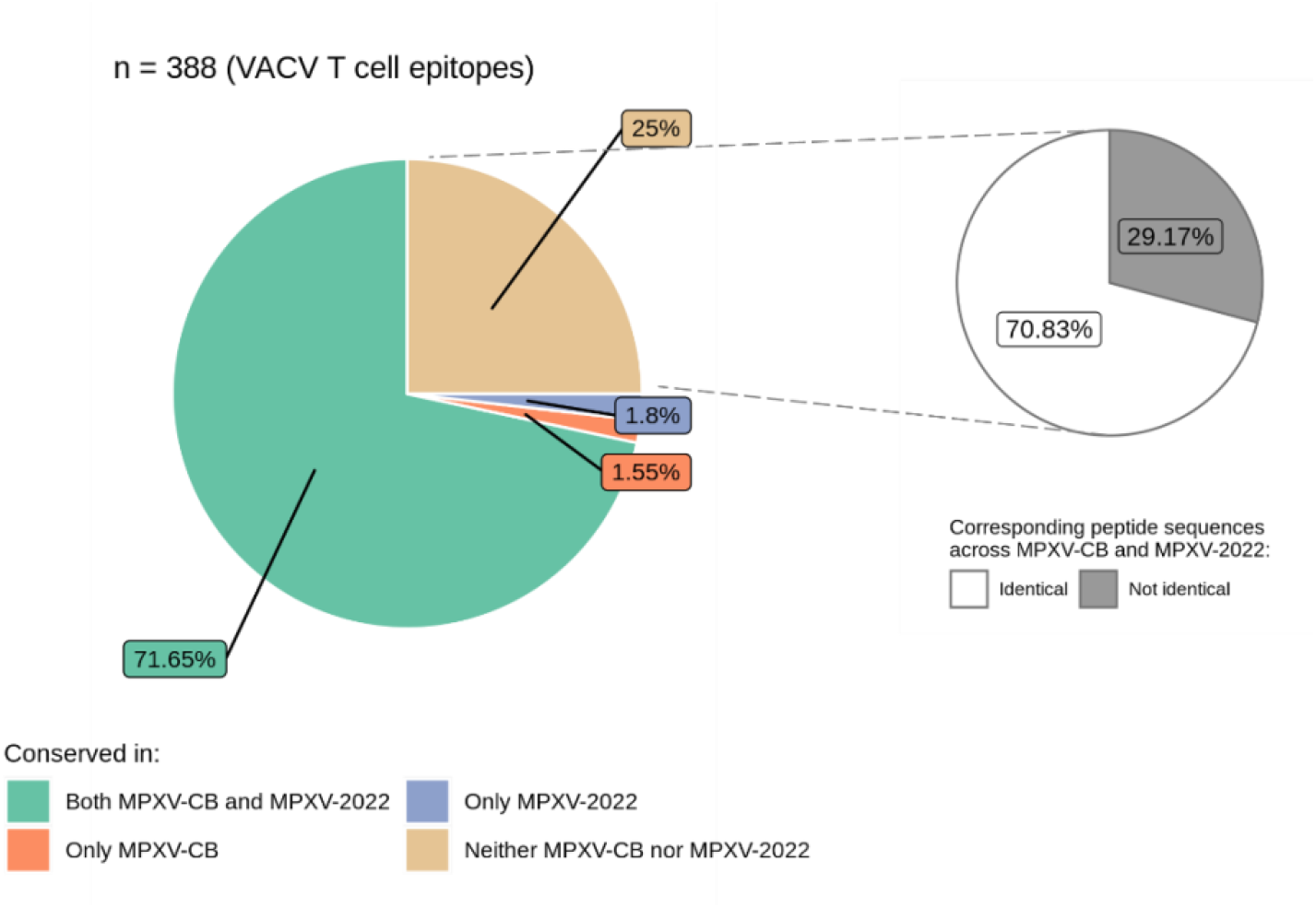
Distribution of the VACV T cell epitopes (n = 388) conserved among MPXV-2022 and MPXV-CB.

## DISCUSSION

The MPXV-2022 outbreak is the first multi-country spread of monkeypox outside Africa [2,3]. The rapid speed at which the outbreak has spread is concerning. While the MPXV-2022 isolates have been associated with the MPXV-WA clade, these differ on average by 50 SNPs from the closest MPXV-WA sequences, which were collected in 2018-2019 [10]. This number of SNPs is surprising when one compares it to the estimated substitution rate (∼1-2 SNPs per genome per year [36,37]) for Orthopoxviruses. These genetic differences raise questions about the evolutionary origin of the additional mutations associated with the 2022 outbreak, as well as their potential effects on viral transmission, infectivity, and immune recognition. Research is being pursued to address these questions.

The current study aimed to investigate to what extent the genetic differences observed in the MPXV-2022 outbreak viruses may be expected to impact immune responses induced by VACV vaccines that are being recommended as interventions to counter monkeypox. This question is further confounded by the recognition that while the effectiveness of the VACV vaccines has been reported for MPXV-CB [4,16], to our knowledge, similar studies are not available for MPXV-WA. Comparing the genetic composition of known targets of VACV-elicited neutralizing antibodies or T cells, either at the epitope level (where available) or the protein level, demonstrated limited genetic variability among the MPXV-2022 sequences. Moreover, the large majority of corresponding genetic variation in MPXV-2022 was commonly observed in MPXV-CB. Based on this, it may be anticipated that the VACV vaccines will elicit similar humoral and cellular immunity against MPXV-2022 as for MPXV-CB. If this is the case, the high efficacy of VACV vaccines that has been reported against MPXV-CB [4,16] may be preserved for MPXV-2022. However, clinical data is required to determine the exact efficacy of VACV vaccines against MPXV-2022.

## DATA AND CODE AVAILABILITY

GISAID acquisition IDs of all monkeypox sequences, IEDB IDs of all epitope data, and all coding scripts (written in the R language) for reproducing the results will be made available online as a GitHub repository.

## ACKNOWLEDGMENTS

We thank all the authors, the originating and submitting laboratories (listed in Supplementary Table 3) for their sequence and metadata shared through GISAID, on which this research is based. We also thank the open sharing of immunological data of VACV by research groups from around the world through the IEDB database. We gratefully acknowledge the contributions of all the researchers, scientists and technical staff involved.

## SUPPLEMENTARY FIGURES AND TABLES

**Supplementary Figure 1.**
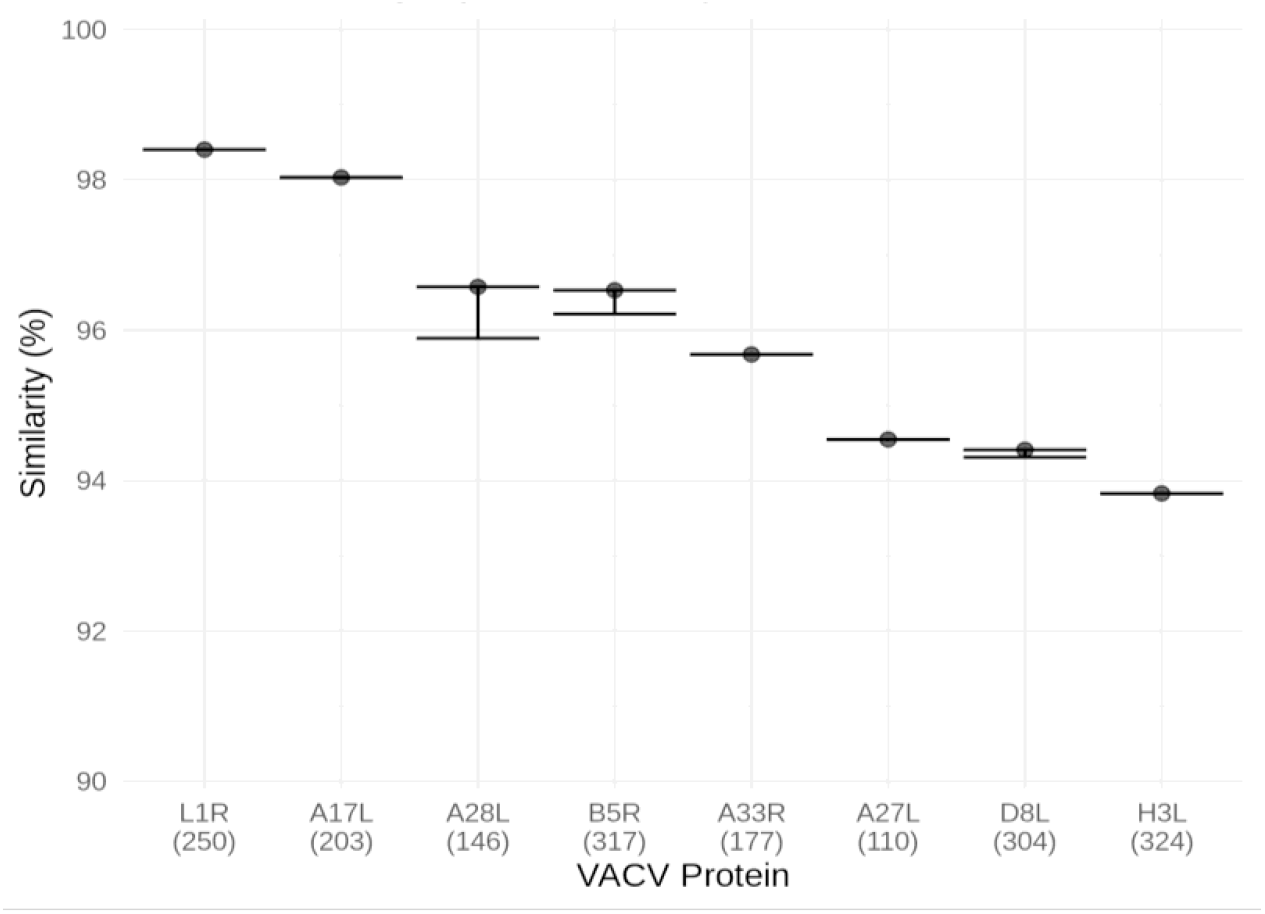
Genetic similarity between VACV proteins targeted by NAbs and their orthologs in 75 MPXV-2022 sequences. A circle represents the median similarity of the VACV protein among the MPXV-2022 sequences, while the lower and upper bars indicate the 5th and 95th percentiles respectively. Length of the VACV proteins is indicated within parentheses on the x-axis.

**Supplementary Figure 2.**
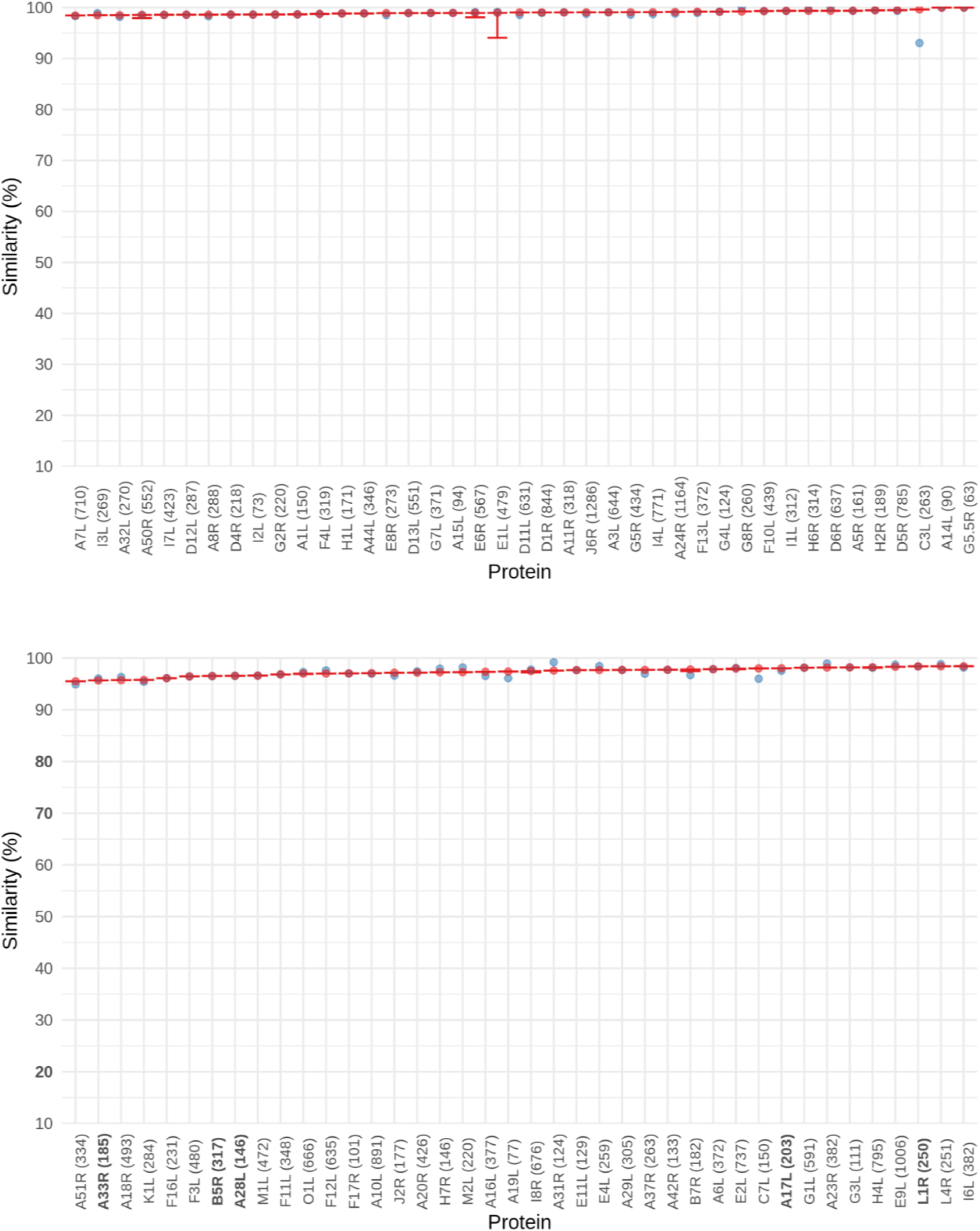
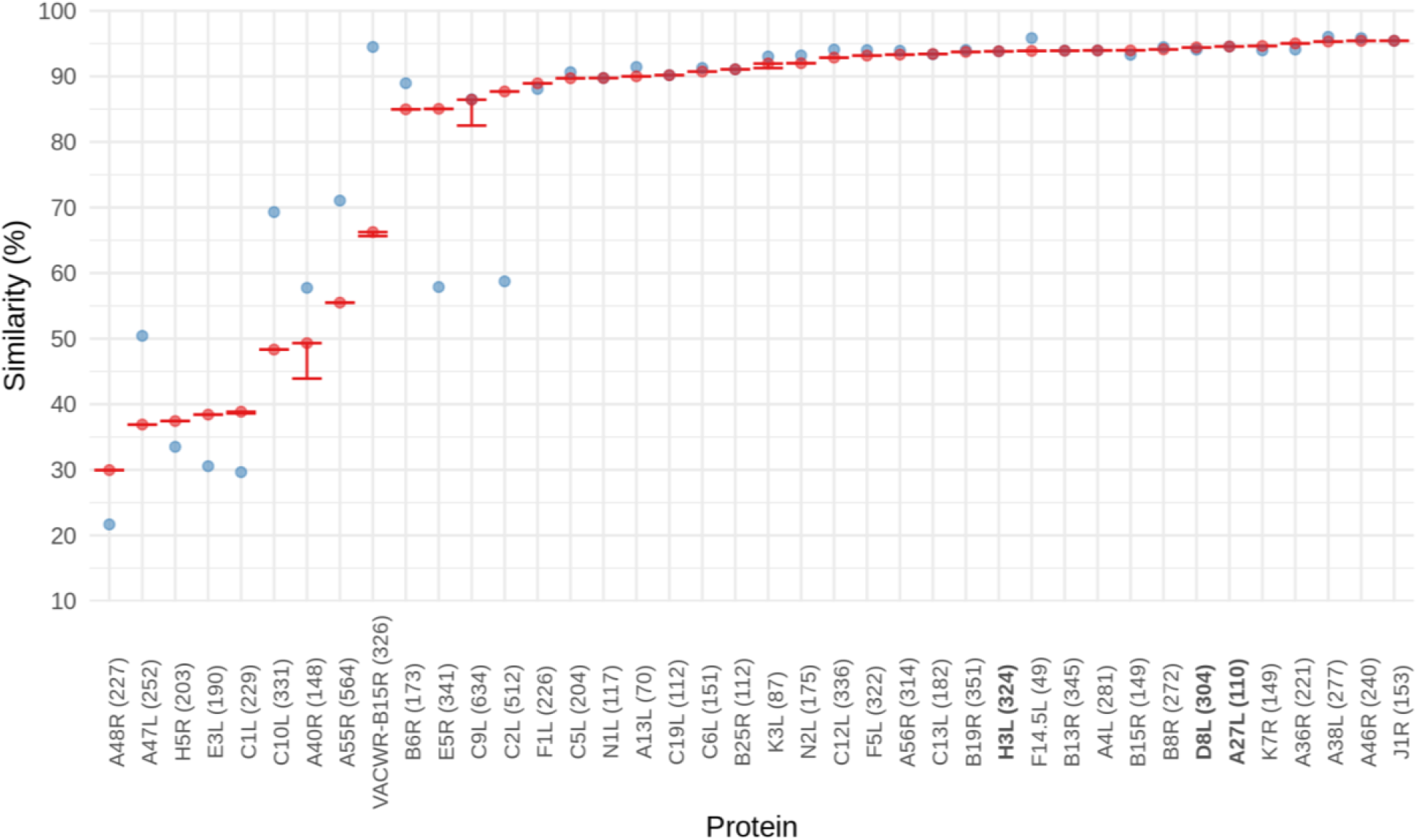
Genetic similarity of 121 VACV-specific proteins with known T cell epitopes among MPXV-2022 (red) and MPXV-CB (blue) orthologs. Variation among the MPXV-2022 sequences is also shown where the lower and upper ends of the error bars indicate the 10th and 90th percentiles respectively. Note that for most proteins the diversity is limited and hence the two ends of the error bars overlap each other. VACV proteins known to be targets of NAbs are indicated in bold. For the nine least similar VACV proteins (A48R, A47L, H5R, E3L, C1L, C10L, A40R, A55R, and VACWR-B15R), there are large spans of indels, which may be a result of partial gene deletions and/or poor quality of the sequence alignment spanning genomic regions that encode these proteins.

**Supplementary Table 1 (included as a separate file)**. List of GISAID Accession IDs of the MPXV-2022 sequences analyzed.

**Supplementary Table 2 (included as a separate file)**. Complete list of experimentally-determined VACV T cell epitopes (n=388) and their conservation in MPXV-2022 and MPXV-CB.

**Supplementary Table 3 (included as a separate file)**. Acknowledgment table (downloaded from GISAID).

