## Supplementary Table 3 for "Vaccinia virus vaccination is expected to elicit highly cross-reactive immunity to the 2022 monkeypox virus"

We gratefully acknowledge the following Authors from the Originating laboratories responsible for obtaining the specimens, as well as the Submitting laboratories where the genome data were generated and shared via GISAID, on which this research is based.

All Submitters of data may be contacted directly via www.gisaid.org

Authors are sorted alphabetically.

| Accession ID | Originating Laboratory | Submitting Laboratory | Authors |
|---|---|---|---|
| EPI_ISL_13052263 | Microbiol Genomics and Bioinformatics, Bundeswehr Institute of Microbiology | Microbiol Genomics and Bioinformatics, Bundeswehr Institute of Microbiology | Antwerpen,M.H., Lang,D., Zange,S., Walter,M.C. and Woelfel,R. |
| EPI_ISL_13052264, EPI_ISL_13052265, EPI_ISL_13052266, EPI_ISL_13052267, EPI_ISL_13052268, EPI_ISL_13052269, EPI_ISL_13052270, EPI_ISL_13052271, EPI_ISL_13052272, EPI_ISL_13052273 | Instituto Nacional de Saude Doutor Ricardo Jorge (INSA) | Instituto Nacional de Saude Doutor Ricardo Jorge (INSA) | Joana Isidro, Vítor Borges, Miguel Pinto, Daniel Sobral, João Dourado Santos, Alexandra Nunes, Verónica Mixão, Rita Ferreira, Daniela Santos, Silvia Duarte, Luís Vieira, Maria José Borrego, Sofia Núncio, Isabel Lopes de Carvalho, Ana Pelerito, Rita Cordeiro, João Paulo Gomes |
| EPI_ISL_13052274 | Laboratory of Virology, University Hospitals of Geneva | Laboratory of Virology, University Hospitals of Geneva | Laubscher,F., Chudzinski,V., Schibler,M., Kaiser,L. and Renzoni,A. |
| EPI_ISL_13052275 | IHAP, VIRAL, Universite de Toulouse, INRAE, ENVT | IHAP, VIRAL, Universite de Toulouse, INRAE, ENVT | Croville,G., Walch,M., Guerin,J.-L., Mansuy,J.-M., Pasquier,C. and Izopet,J. |
| EPI_ISL_13052277 | Public Health Virology, Erasmus Medical Centre | Public Health Virology, Erasmus Medical Centre | Oude Munnink,B.B., Boter,M., Wellers,B., Molenkamp,R., Sikkema,R.S. and Koopmans,M. |
| EPI_ISL_13052278 | Research and Evaluation, UKHSA | Research and Evaluation, UKHSA | Osman,K.L., Lewandowski,K.S., Pullan,S.T., Carter,D.P., Crook,J.M., Vipond,R. and Chand,M. |
| EPI_ISL_13052279, EPI_ISL_13052280, EPI_ISL_13052281 | Research and Evaluation, UKHSA | Research and Evaluation, UKHSA | Osman,K.L., Lewandowski,K.S., Carter,D.P., Crook,J.M., Pullan,S.T., Vipond,R. and Chand,M. |
| EPI_ISL_13052282 | Microbiology, Immunology and Transplantation, KU Leuven, Rega Institute | Microbiology, Immunology and Transplantation, KU Leuven, Rega Institute | Vanmechelen,B., Wawina-Bokalanga,T., Logist,A.-S., Sinnesael,R., Ysebaert,L., Verlinden,J., Bloemen,M. and Maes,P. |
| EPI_ISL_13052283 | Microbiology, Immunology and Transplantation, KU Leuven, Rega Institute | Microbiology, Immunology and Transplantation, KU Leuven, Rega Institute | Wawina-Bokalanga,T., Vanmechelen,B., Logist,A.-S., Sinnesael,R., Ysebaert,L., Verlinden,J., Bloemen,M. and Maes,P. |
| EPI_ISL_13052284 | Microbiology, Hospital Universitari Germans Trias i Pujol | Microbiology, Hospital Universitari Germans Trias i Pujol | Martinez-Puchol,S., Coello,A., Bordoy,A.E., Soler,L., Panisello,D., Gonzalez-Gomez,S., Clara,G., Paris de Leon,A., Not,A., Hernandez,A., Bofill-Mas,S., Saludes,V., Blanco,I., Martro,E. and Cardona,P.-J. |
| EPI_ISL_13052285 | Laboratory of Virology, University Hospitals of Geneva | Laboratory of Virology, University Hospitals of Geneva | Laubscher,F., Schibler,M., Kaiser,L. and Renzoni,A. |
| EPI_ISL_13052286 | Department of Biomedical and Clinical Sciences, University of Milan | Department of Biomedical and Clinical Sciences, University of Milan | Lai,A., Bergna,A., Della Ventura,C., Tarkowski,M., Riva,A., Moschese,D., Rizzardini,G., Antinori,S. and Zehender,G. |
| EPI_ISL_13052287 | Virology, GENomique EPIdemiologique des maladies Infectieuses | Virology, GENomique EPIdemiologique des maladies Infectieuses | unknown |
| EPI_ISL_13052288 | Department of Health, Utah Public Health Laboratory | Department of Health, Utah Public Health Laboratory | Young,E.L., Hergert,J. and Oakeson,K.F. |
| EPI_ISL_13052289 | Division of High-Consequence Pathogens and Pathology, Centers for Disease Control and Prevention | Division of High-Consequence Pathogens and Pathology, Centers for Disease Control and Prevention | Gigante,C.M., Smole,S., Seabolt,M.H., Wilkins,K., McCollum,A., Hutson,C., Davidson,W., Rao,A., Brown,C. and Li,Y. |
| EPI_ISL_13052290 | Laboratory for Diagnostics of Zoonoses and WHO Centre, Institute of Microbiology and Immunology, Faculty of Medicine, University of Ljubljana | Laboratory for Diagnostics of Zoonoses and WHO Centre, Institute of Microbiology and Immunology, Faculty of Medicine, University of Ljubljana | Zakotnik,S., Vlaj,D., Suljic,A., Zorec,T.M., Korva,M., Poljak,M. and Avsic Zupanc,T. |
| EPI_ISL_13052291 | Laboratory for Diagnostics of Zoonoses and WHO Centre, Institute of Microbiology and Immunology, Faculty of Medicine, University of Ljubljana | Laboratory for Diagnostics of Zoonoses and WHO Centre, Institute of Microbiology and Immunology, Faculty of Medicine, University of Ljubljana | Zakotnik,S., Vlaj,D., Suljic,A., Zorec,T.M., Korva,M. and Avsic Zupanc,T. |
| EPI_ISL_13052292 | Victorian Infectious Diseases Reference Laboratory, Doherty Institute | Victorian Infectious Diseases Reference Laboratory, Doherty Institute | Hammerschlag,Y., MacLeod,G., Papadakis,G., Adan-Sanchez,A., Druce,J.D., Williamson,D.A., Cheng,A.C. and McMahon,J.H. |
| EPI_ISL_13052293, EPI_ISL_13052294 | Centre for Biological Threats, Highly Pathogenic Viruses, Robert Koch Institute | Centre for Biological Threats, Highly Pathogenic Viruses, Robert Koch Institute | Brinkmann,A., Kohl,C., Uddin,S., Pape,K., Schrick,L., Michel,J., Schaade,L. and Nitsche,A. |
| EPI_ISL_13052295 | SC (UCO) Igiene e Sanità Pubblica, ASUGI, Trieste | Genomics and Epigenomics, AREA Science Park | Licastro,D., DeGasperi,M., Negri,C., Piscianz,E., Koncan,R., Dal Monego,S., Segat,L. and D'Agaro,P. |
| EPI_ISL_13056892, EPI_ISL_13056893, EPI_ISL_13056894, EPI_ISL_13056895, EPI_ISL_13056896, EPI_ISL_13056897, EPI_ISL_13056898, EPI_ISL_13056899, EPI_ISL_13056900, EPI_ISL_13056901, EPI_ISL_13056902, EPI_ISL_13056903, EPI_ISL_13056904, EPI_ISL_13056905, EPI_ISL_13056906, EPI_ISL_13056907, EPI_ISL_13056908, EPI_ISL_13056909 | | | |
| see above | Instituto Nacional de Saude Doutor Ricardo Jorge (INSA) | Instituto Nacional de Saude Doutor Ricardo Jorge (INSA) | Joana Isidro, Vítor Borges, Miguel Pinto, Daniel Sobral, João Dourado Santos, Alexandra Nunes, Verónica Mixão, Rita Ferreira, Daniela Santos, Silvia Duarte, Luís Vieira, Maria José Borrego, Sofia Núncio, Isabel Lopes de Carvalho, Ana Pelerito, Rita Cordeiro, João Paulo Gomes |
| EPI_ISL_13056910 | Biochemistry and Molecular Genetics, Israel Institute for Biological Research | Biochemistry and Molecular Genetics, Israel Institute for Biological Research | Israeli,O., Guedj-Dana,Y., Lazar,S., Shifman,O., Erez,N., Weiss,S., Paran,N., Israely,T., Schuster,O., Zvi,A., Beth-Din,A. and Cohen Gihon,I. |
| EPI_ISL_13089461 | Hospital General Universitario Gregorio Marañón | Hospital General Universitario Gregorio Marañón | Sergio Buenestado Serrano, Rosalía Palomino Cabrera, Daniel Peñas Utrilla, Jorge Rodríguez-Grande, Laura Pérez-Lago, Cristina Rodriguez-Grande, Marta Herranz Martin, Julia Suárez, Pilar Catalán, Patricia Muñoz, Darío García de Viedma |
| EPI_ISL_13094227 | Division of High-Consequence Pathogens and Pathology, Centers for Disease Control and Prevention | Division of High-Consequence Pathogens and Pathology, Centers for Disease Control and Prevention | Gigante,C.M., Lee,P., Seabolt,M.H., Wilkins,K., McCollum,A., Hutson,C., Davidson,W., Rao,A., Mendoza,R. and Li,Y. |
| EPI_ISL_13096615 | Division of High-Consequence Pathogens and Pathology, Centers for Disease Control and Prevention | Division of High-Consequence Pathogens and Pathology, Centers for Disease Control and Prevention | Gigante,C.M., Griffin-Thomas,L.A., Seabolt,M.H., Wilkins,K., McCollum,A., Hutson,C., Davidson,W., Rao,A., Crain,J. and Li,Y. |
| EPI_ISL_13100618 | Division of High-Consequence Pathogens and Pathology, Centers for Disease Control and Prevention | Division of High-Consequence Pathogens and Pathology, Centers for Disease Control and Prevention | Gigante,C.M., Ventura,J., Seabolt,M.H., Wilkins,K., McCollum,A., Hutson,C., Davidson,W., Rao,A., Nash,J. and Li,Y. |
| EPI_ISL_13100619 | Division of High-Consequence Pathogens and Pathology, Centers for Disease Control and Prevention | Division of High-Consequence Pathogens and Pathology, Centers for Disease Control and Prevention | Gigante,C.M., Lee,P., Seabolt,M.H., Wilkins,K., McCollum,A., Hutson,C., Davidson,W., Rao,A., Mendoza,R. and Li,Y. |
| EPI_ISL_13100620 | Division of High-Consequence Pathogens and Pathology, Centers for Disease Control and Prevention | Division of High-Consequence Pathogens and Pathology, Centers for Disease Control and Prevention | Gigante,C.M., Atkinson,A., Seabolt,M.H., Wilkins,K., McCollum,A., Hutson,C., Davidson,W., Rao,A., Murray,J. and Li,Y. |
| EPI_ISL_13100621 | Division of High-Consequence Pathogens and Pathology, Centers for Disease Control and Prevention | Division of High-Consequence Pathogens and Pathology, Centers for Disease Control and Prevention | Gigante,C.M., Stringer,J., Seabolt,M.H., Wilkins,K., McCollum,A., Hutson,C., Davidson,W., Rao,A., Schulte,J. and Li,Y. |
| EPI_ISL_13100622 | Division of High-Consequence Pathogens and Pathology, Centers for Disease Control and Prevention | Division of High-Consequence Pathogens and Pathology, Centers for Disease Control and Prevention | Gigante,C.M., Myers,R., Seabolt,M.H., Wilkins,K., McCollum,A., Hutson,C., Davidson,W., Rao,A., Blythe,D. and Li,Y. |

| | | | |
|---|---|---|---|
| EPI_ISL_13100719 | Division of High-Consequence Pathogens and Pathology, Centers for Disease Control and Prevention | Division of High-Consequence Pathogens and Pathology, Centers for Disease Control and Prevention | Gigante,C.M., Ventura,J., Seabolt,M.H., Wilkins,K., McCollum,A., Hutson,C., Davidson,W., Rao,A., Nash,J. and Li,Y. |
| EPI_ISL_13106454 | Hospital General Universitario Gregorio Marañón | Hospital General Universitario Gregorio Marañón | Sergio Buenestado Serrano, Rosalía Palomino Cabrera, Daniel Peñas Utrilla, Jorge Rodríguez-Grande, Pedro Sola Campoy, Laura Pérez-Lago, Cristina Rodriguez-Grande, Marta Herranz Martin, Julia Suárez, Pilar Catalán, Patricia Muñoz, Darío García de Viedma |
| EPI_ISL_13117291 | Centre for Biological Threats, Highly Pathogenic Viruses, Robert Koch Institute | Centre for Biological Threats, Highly Pathogenic Viruses, Robert Koch Institute | Brinkmann,A., Kohl,C., Uddin,S., Pape,K., Schrick,L., Michel,J., Jessen,H., Schaade,L. and Michel,A. |
| EPI_ISL_13117292, EPI_ISL_13117293 | Centre for Biological Threats, Highly Pathogenic Viruses, Robert Koch Institute | Centre for Biological Threats, Highly Pathogenic Viruses, Robert Koch Institute | Brinkmann,A., Kohl,C., Uddin,S., Pape,K., Schrick,L., Michel,J., Stocker,H., Schaade,L. and Nitsche,A. |
| EPI_ISL_13117294, EPI_ISL_13117295, EPI_ISL_13117296, EPI_ISL_13117297, EPI_ISL_13117298 | Centre for Biological Threats, Highly Pathogenic Viruses, Robert Koch Institute | Centre for Biological Threats, Highly Pathogenic Viruses, Robert Koch Institute | Brinkmann,A., Kohl,C., Uddin,S., Pape,K., Schrick,L., Michel,J., Schaade,L. and Nitsche,A. |
| EPI_ISL_13148263, EPI_ISL_13148264, EPI_ISL_13148265 | Centre for Biological Threats, Highly Pathogenic Viruses, Robert Koch Institute | Centre for Biological Threats, Highly Pathogenic Viruses, Robert Koch Institute | Brinkmann,A., Kohl,C., Uddin,S., Pape,K., Schrick,L., Michel,J., Jessen,H., Schaade,L. and Nitsche,A. |
| EPI_ISL_13148266, EPI_ISL_13148267, EPI_ISL_13148268, EPI_ISL_13148269 | Centre for Biological Threats, Highly Pathogenic Viruses, Robert Koch Institute | Centre for Biological Threats, Highly Pathogenic Viruses, Robert Koch Institute | Brinkmann,A., Kohl,C., Uddin,S., Pape,K., Schrick,L., Michel,J., Stocker,H., Schaade,L. and Nitsche,A. |
| EPI_ISL_13148270, EPI_ISL_13148271, EPI_ISL_13148272, EPI_ISL_13148273, EPI_ISL_13148274, EPI_ISL_13148275, EPI_ISL_13148276 | Centre for Biological Threats, Highly Pathogenic Viruses, Robert Koch Institute | Centre for Biological Threats, Highly Pathogenic Viruses, Robert Koch Institute | Brinkmann,A., Kohl,C., Uddin,S., Pape,K., Schrick,L., Michel,J., Schaade,L. and Nitsche,A. |
